# YME1L1 is Dispensable for T Lymphocyte Activation Despite its Upregulation and Activity

**DOI:** 10.64898/2026.03.16.712079

**Authors:** Gonçalo Malpica, Mariana Joaquim, Ricardo Silva Machado, João Fernandes, Michael J. Hall, Gabriel Martins, Vanessa A. Morais, Marc Veldhoen

## Abstract

Mitochondrial dynamics are critical for T cell activation, differentiation, and survival. The inner mitochondrial membrane ATP-dependent metalloprotease YME1L1 regulates proteostasis and the processing of optic atrophy protein 1 (OPA1), thereby shaping mitochondrial cristae architecture and respiratory function in many cell types. Whether YME1L1 fulfils similar roles in lymphocytes remains unknown. Here, we examined YME1L1 function in T cells using conditional knockout mice lacking YME1L1 in lymphocytes (YME1L1^ΔTB^). YME1L1 expression increased upon T cell activation, yet its absence did not alter thymic development, peripheral T cell homeostasis, or the proportions of naïve, memory, and regulatory subsets. T cell activation and proliferation in response to anti-CD3ε stimulation were also unaffected. Mitochondrial parameters such as mass, membrane potential, and reactive oxygen species production, were largely preserved, with only modest, transient increases in oxidative stress detected in CD4 T cells lacking YME1L1. Electron microscopy revealed no major changes in mitochondrial size or roundness but showed increased cristae branching and reduced tortuosity, indicating subtle alterations in ultrastructure. Additionally, γδ T cells in YME1L1^ΔTB^ mice exhibited a mild shift toward interferon-γ–producing phenotypes at the expense of interleukin-17–producing subsets. Collectively, our data indicate that YME1L1, despite its requirement for OPA1 cleavage, is dispensable for T cell development and acute activation but may contribute to fine-tune mitochondrial architecture and γδ T cell effector programming. These findings highlight cell-type–specific redundancies in mitochondrial quality control and underscore the value of negative data in refining the understanding of mitochondrial regulation in immune cells.

## Introduction

T lymphocytes are reported to be highly dependent on mitochondrial remodelling to support their functional states, which needs to change quickly upon recognising their cognate antigen [1]. During their activation, T cells undergo profound metabolic reprogramming, shifting from a quiescent, oxidative state to one that integrates increased glycolysis with mitochondrial oxidative phosphorylation to sustain rapid proliferation and effector functions [2, 3]. Memory T cells, in contrast, rely more heavily on oxidative phosphorylation and fatty acid oxidation, processes that are also tightly linked to mitochondrial dynamics and cristae organisation, with some tissue resident T cells maintaining their mitochondria in an altered state resembling their poised activity [4].

Mitochondria are highly dynamic organelles, recognised for their role in ATP synthesis and as the energetic hub of cells. Beyond their bioenergetic role, they are signalling platforms that contribute to diverse cellular processes, in particular calcium buffering, production of reactive oxygen species, and programmed cell death [5–7]. Furthermore, they have important roles in anabolic processes to generate metabolites required for lipids, nucleotide and protein synthesis [8]. The maintenance of a healthy mitochondrial network depends on a balance between their biogenesis, breakdown via mitophagy, and the dynamic processes of fusion and fission. These events are coordinated by GTPases that remodel mitochondrial membranes and by factors that regulate cristae organisation, ensuring efficient oxidative phosphorylation and thereby cellular function [9].

A central player in these processes is the dynamin-like GTPase optic atrophy protein-1 (OPA1), which resides in the inner mitochondrial membrane (IMM) and is critical for both inner membrane fusion and cristae architecture. The IMM can fold in on itself to form cristae that greatly expand surface area and create specialized loci for distinct functions [10]. By controlling the cristae shape, OPA1 influences the assembly of respiratory chain complexes, mitochondrial ATP generating efficiency, and thereby several processes like regulation of apoptosis [9, 11–13]. The activity and stability of OPA1 itself is, in turn, tightly regulated by mitochondrial proteases, a process that results in different forms of OPA1. Long forms of OPA1 (L-OPA1) are proteolytically cleaved by two IMM proteases to generate S-OPA1 forms [14].

Among the proteases that target OPA1, the ATP-dependent zinc metalloprotease YME1L1, also located in the IMM, has emerged as a key determinant of proteostasis [15, 16]. YME1L1 degrades misfolded or damaged proteins and processes substrates such as OPA1, thereby modulating cristae structure, mitochondrial dynamics, and respiratory chain assembly [11, 17]. Through these actions, YME1L1 safeguards mitochondrial homeostasis and influences cellular metabolism and survival [17, 18]. Modifications that alter mitochondrial dynamics will change its function, which can result in cellular dysregulation [19]. Regulators such as OPA1 have been shown to be indispensable for maintaining mitochondrial integrity and function in lymphocytes [20], underscoring the importance of inner membrane structure in immune responses [21]. The ratio of L- and S-OPA1 governs IMM fusion events and cristae organization, regulated by the cleaving of L-OPA1 by only two enzymes, YME1L1 and OMA1 [12, 22]. Whether YME1L1 fulfils such functions in T lymphocytes has not been addressed.

Given that YME1L1 directly processes OPA1 and controls the turnover of multiple inner membrane proteins, it would be expected to play a critical role in mitochondrial dynamics and cell function. Although the absence of YME1L1 results in the stabilisation of one of the L-OPA1 forms, only minor differences in mitochondrial fission and fusion have been reported in mouse embryonic fibroblasts lacking YME1L1, which show an increase in mitochondrial fragmentation [23]. This observation is in line with reports of the role of YME1L1 in early porcine oocytes, in which increased reactive oxygen species, reduced mitochondrial membrane potential and ATP production were reported as a result of YME1L1 knockdown [24]. In non-small cell lung and osteosarcoma cancer cells, YME1L1 loss also leads to a decrease in mitochondrial membrane potential, higher oxidative stress and lower ATP production, as well as lower cell size and loss of proliferative capacity [25, 26]. Furthermore, in mouse embryonic fibroblasts and patient derived fibroblasts, as well as mouse neurons, YME1L1 loss was associated with fragmented mitochondria [11, 17, 23, 27–31]. In addition, in human embryonic kidney cells this was accompanied by cristae disorganization and loss of respiratory capacity [28, 32]. Since T cell development, activation, and survival have a high dependency on mitochondrial activity we hypothesised a role for YME1L1 in T cell biology.

## Results

### YME1L1 levels increase in activated T cells

To investigate a potential role of YME1L1 in T lymphocytes, we assessed its protein levels in B and CD4 and CD8 T lymphocytes under steady state conditions. YME1L1 protein levels were similar in lymphocyte subsets (Fig 1a-b). The rapid proliferation of T cells is accompanied by protein synthesis and metabolic switching of the cell metabolism programme [33]. In agreement with this, upon *in vivo* activation, using intraperitoneal injection of anti-CD3, the level of YME1L1 protein is increased, peaking within the first 24 hours in both CD4 and CD8 T cells (Fig. 1c-e). Of interest, we did not observe a similar trend in B cells obtained from the same mice injected with anti-CD3, despite their heightened activation state (Fig. 1f).

**Figure 1.**
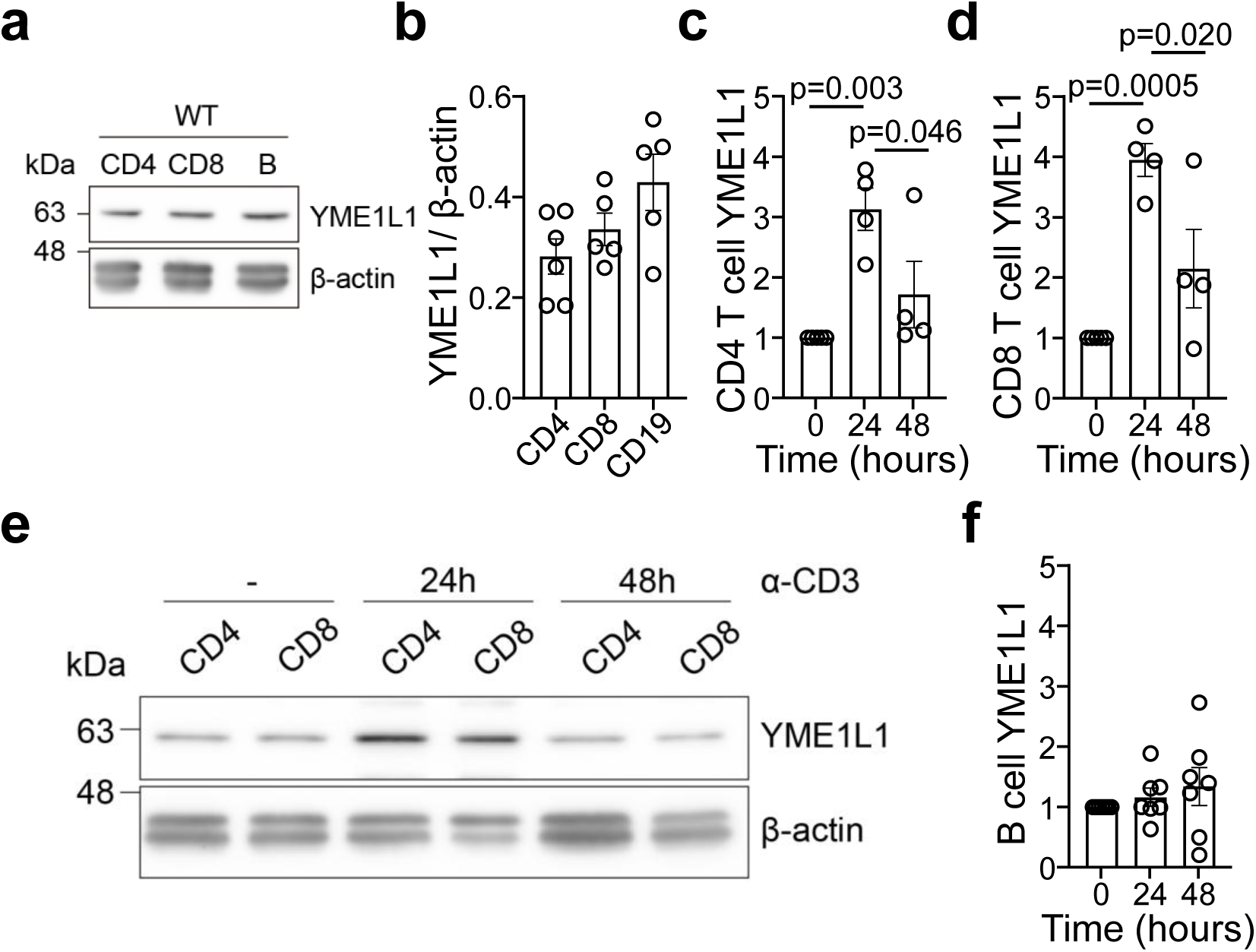
YME1L1 protein levels increase upon T cell activation. CD4 and CD8 T cells or B cells were harvested from C57BL6/J mice at steady state or upon indicated time after intraperitoneal administration of anti-CD3ε. **a**) Representative Western Blot with levels of YME1L1 and β-actin in indicate cell types. **b**) YME1L1 protein level relative to β-actin assessed in indicated lymphocyte populations. **c-f**) Fold increase of YME1L1/β-actin protein level from steady state upon indicated time after anti-CD3ε administration in c) CD4 T cells, d) CD8 T cells, e) representative Western Blot of CD4 and CD8 T cells, f) B cells (n=4). Statistical analysis: One-way ANOVA, Turkeýs multiple comparison test.

### YME1L1 deletion does not affect peripheral T cell proportions and levels

To ascertain a role for YME1L1 in T cells, we obtained the YME1L1 conditional knock out mice [17]. These were crossed with the Rag1-Cre line to obtain lymphocyte specific deletion [34]. In these mice, referred to as YME1L1^ΔTB^, YME1L1 protein is absent in both B and T cells (Fig. 2a). In the absence of YME1L1, the proportion and level of total B lymphocytes is similar in the spleens from YME1L1^ΔTB^ and YME1L1^fl/fl^ controls (Fig. 2b). Furthermore, this was also found for the numbers and proportions of both CD4 and CD8 T cells (Fig. 2c-d). A more detailed assessment of naïve, memory and regulatory (Treg) CD4 T cells as well as naïve and memory CD8 T cells did not reveal any proportional differences between both genotypes (Fig. 2e-g). Despite the increased YME1L1 protein levels upon T cell activation, the presence and distribution of T cell subsets was not altered in the absence of YME1L1.

**Figure 2.**
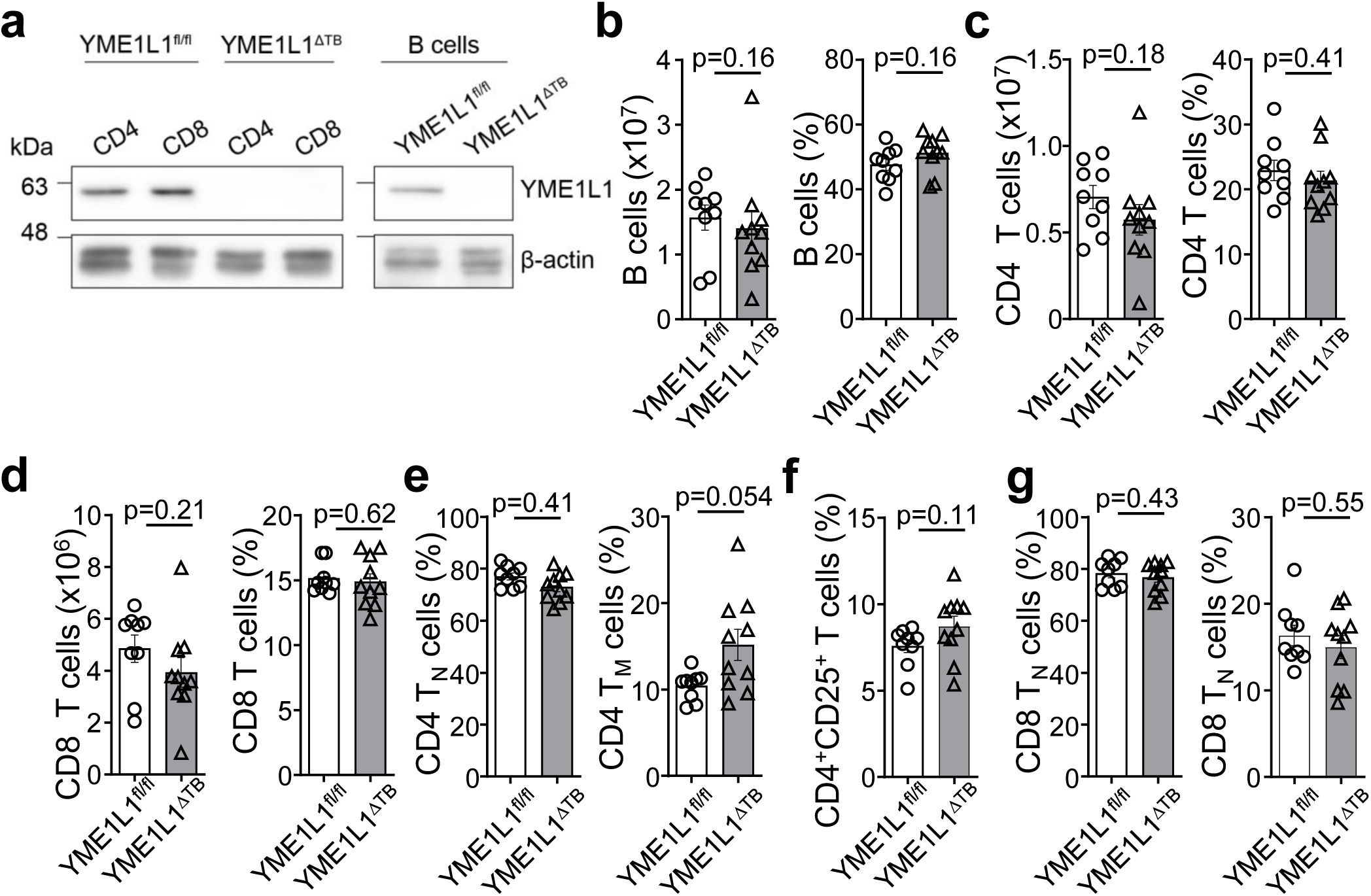
YME1L1-deficiency does not alter proportion and presence of peripheral lymphocyte subsets. Proportions of indicated lymphocyte subsets were assessed in spleens harvested from YME1L1^f/f^ littermate controls and YME1L1^ΔTB^ animals. **a**) Representative Western blot analysis of YME1L1 and β-actin proteins in CD4 and CD8 T cells and B cells from C57BL6/J mice with indicated genotype. **b-g**) Total cell number and proportion of total lymphocytes was assessed of b) B lymphocytes, c) CD4 T cells, d) CD8 T cells; e) proportions of naive and memory CD4 T cells and f) proportion of Treg cells of total CD4 T cells: g) proportion of naïve and memory CD8 T cells of total CD8 T cells (n=9, 10). Statistical analysis: Mann-Whitney tests.

### YME1L1 deletion does not affect developing T cell numbers and proportions

T cells undergo stringent selection in the thymus where useful TCRs are selected and potential autoreactive TCRs are removed from the repertoire. These processes of positive and negative selection are a bottle neck for T cell development and involve TCR signalling events as well as T cell proliferation. To understand if the excision of *Yme1l1* has an impact on T cell development, we assessed the population of T cells upon arrival in the thymus, during selection and the resulting CD4 and CD8 subsets that become part of the circulating T cell populations.

T cell progenitors found in the double negative (DN) population were similar in number and proportions between YME1L1^ΔTB^ and as YME1L1^fl/fl^ controls (Fig. 3a). The large population of double positive (DP) cells did not show signs of alterations such as observed during a T cell developmental block (Fig. 3b). A more detailed analysis of T cell precursors undergoing TCR reengagement and selection, sequentially identified as CD69^-^TCR^lo^, CD69^+^TCR^lo,^ CD69^+^TCR^hi^, and CD69^-^TCR^hi^, showed indistinguishable proportions between YME1L1-deficient T cells and littermate controls (Fig. 3c-f). Furthermore, the resulting numbers and proportions of single positive (SP) CD4 and CD8 T cells was not altered (Fig. 3g-h).

**Figure 3.**
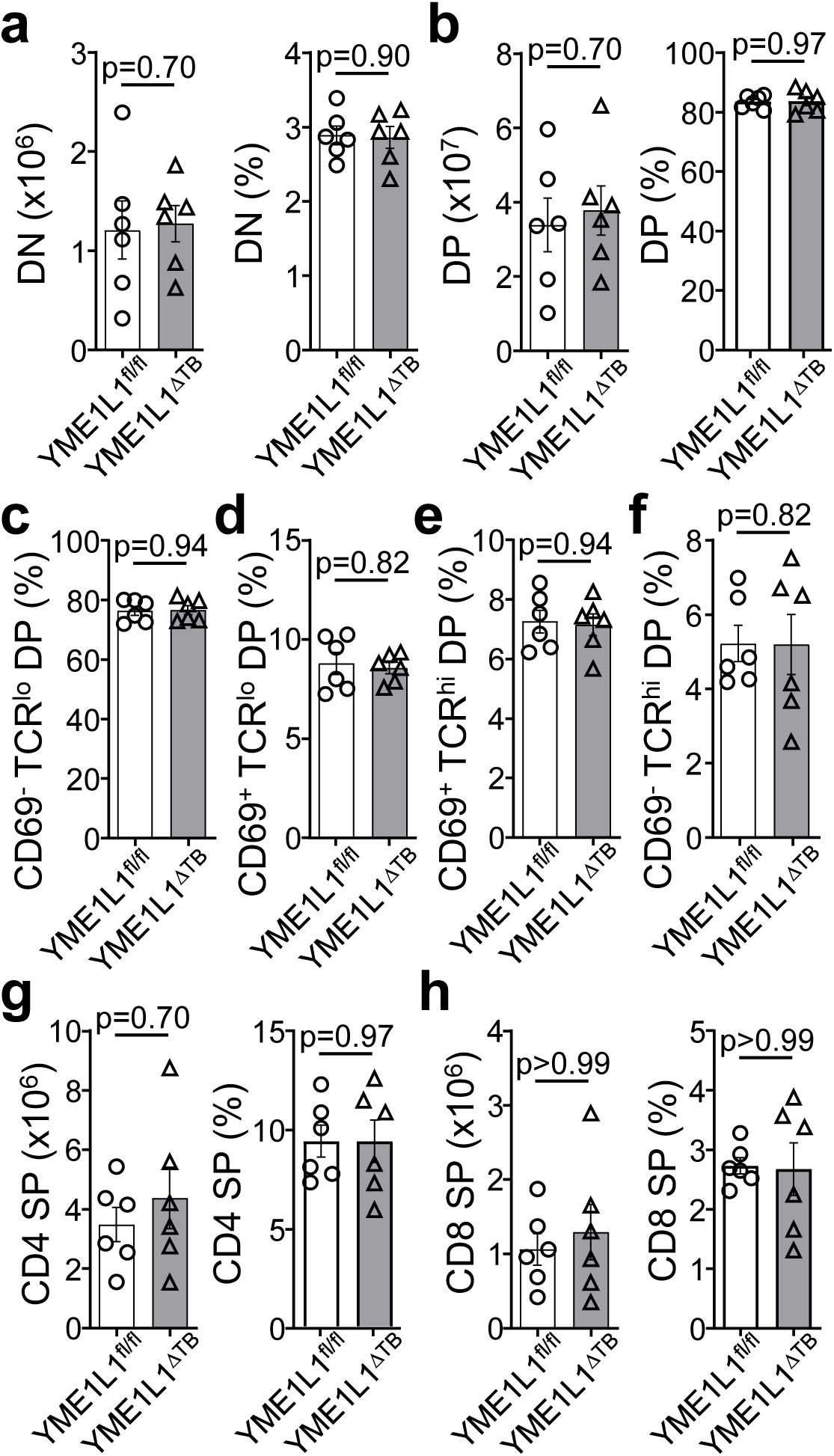
YME1L1-deficiency does not alter proportion and presence of thymic lymphocyte subsets. Proportions of indicated lymphocyte subsets were assessed in the thymus harvested from YME1L1^f/f^ littermate controls and YME1L1^ΔTB^ animals. **a-h**) Total cell number and proportion of total thymocytes was assessed of a) double negative (DN) thymocytes, b) double positive (DP) thymocytes; proportions of DP thymocytes during T cell receptor (TCR) rearrangement and selection c) CD69^-^TCR^lo^, d) CD69^+^TCR^lo^, e) CD69^+^TCR^hi^, and f) CD69^-^TCR^hi^; g) number and proportion of single positive (SP) CD4 T cells and h) number and proportion of SP CD8 T cell of total thymocytes (n=6). Statistical analysis: Mann-Whitney tests.

### YME1L1 deletion does not affect γδ T cell numbers and proportions

In addition to αβ T cells, γδ T cells develop in the thymus and migrate to the periphery, circulating through secondary lymphoid organs and homing to specific tissues. Although these cells show overlaps with αβ T cells, their TCR repertoire is less diverse, and functional properties are in part determined in the thymus. Largely two distinct subsets are recognised - γδ T cells able to produce interferon (IFN)-γ or those able to produce interleukin (IL)-17, amongst other factors [35]. The total number and proportions of γδ T cells in the spleen were not altered in the absence of YME1L1 (Fig. 4a). Although not statistically different, a small tendency towards more γδ T cells able to produce IFN-γ at a cost of those able to produce IL-17 was observed (Fig. 4b-c). To investigate this in more detail, we assessed the thymus. Here, total numbers and proportions of γδ T cells were also similar between both genotypes (Fig. 4d). However, the proportion of cell able to produce IFN-γ was increased, with a decrease in γδ T cells able to make IL-17 in the YME1L1^ΔTB^ compared to their YME1L1^fl/fl^ litter mate controls (Fig. 4e-f).

**Figure 4.**
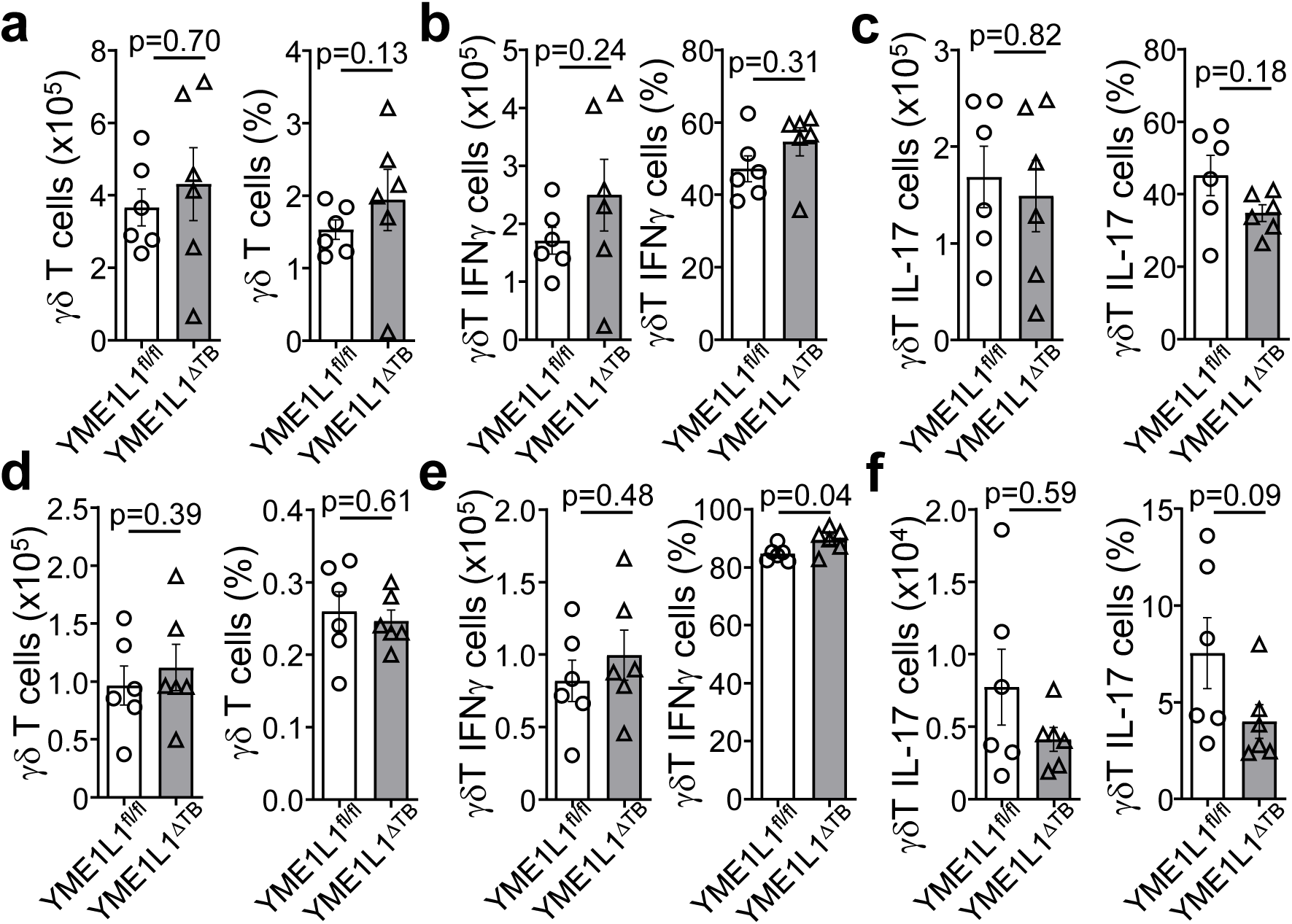
YME1L1-deficiency has limited impact on proportion and presence of thymic and peripheral. γδ**T cell subsets.** Proportions of indicated γδT cell subsets were assessed in the thymus and spleens harvested from YME1L1^f/f^ littermate controls and YME1L1^ΔTB^ animals. **a-c**) Splenic numbers and proportions of a) total γδT cells, b) interferon (IFN)-γ producing γδT cells, and c) interleukin (IL)-17 producing γδT cells. **d-f**) Thymic numbers and proportions of d) total γδT cells, e) interferon (IFN)-γ producing γδT cells, and f) interleukin (IL)-17 producing γδT cells (n=6). Statistical analysis: Mann-Whitney tests.

### T cell proliferation and activation are not impacted by the absence of YME1L1

The absence of YME1L1 or its target OPA1 has been linked to increased susceptibility to apoptosis and reduced proliferation [25, 32, 36]. T cells undergo a rapid and robust proliferative burst upon TCR ligation, accompanied by initial migration inhibition in the secondary lymphoid organs via the expression of the lectin CD69. We mimicked TCR engagement via the intraperitoneal administration of anti-CD3ε antibodies, causing system wide polyclonal T cell activation [3].

We compared the presence of the proliferation marker Ki67 upon 0, 24, 48 or 72 hours of anti-CD3ε administration in YME1L1^ΔTB^ animals and compared these to their litter mate controls. T cells of both genotypes proliferated similarly, increasing Ki67 levels at 24 hours and peaking at 48 hours post activation, after which proliferation reduced. This was similar for CD4 and CD8 T cells (Fig. 5a-b). Furthermore, the lectin CD69 was rapidly expressed at 24 hours post activation on both CD4 and CD8 T cells, after which it was downregulated. This was similar in the absence or presence of YME1L1 (Fig. 5c-d). Collectively, our data suggest that YME1L1 is not required for T cell development, maintenance or proliferation.

**Figure 5.**
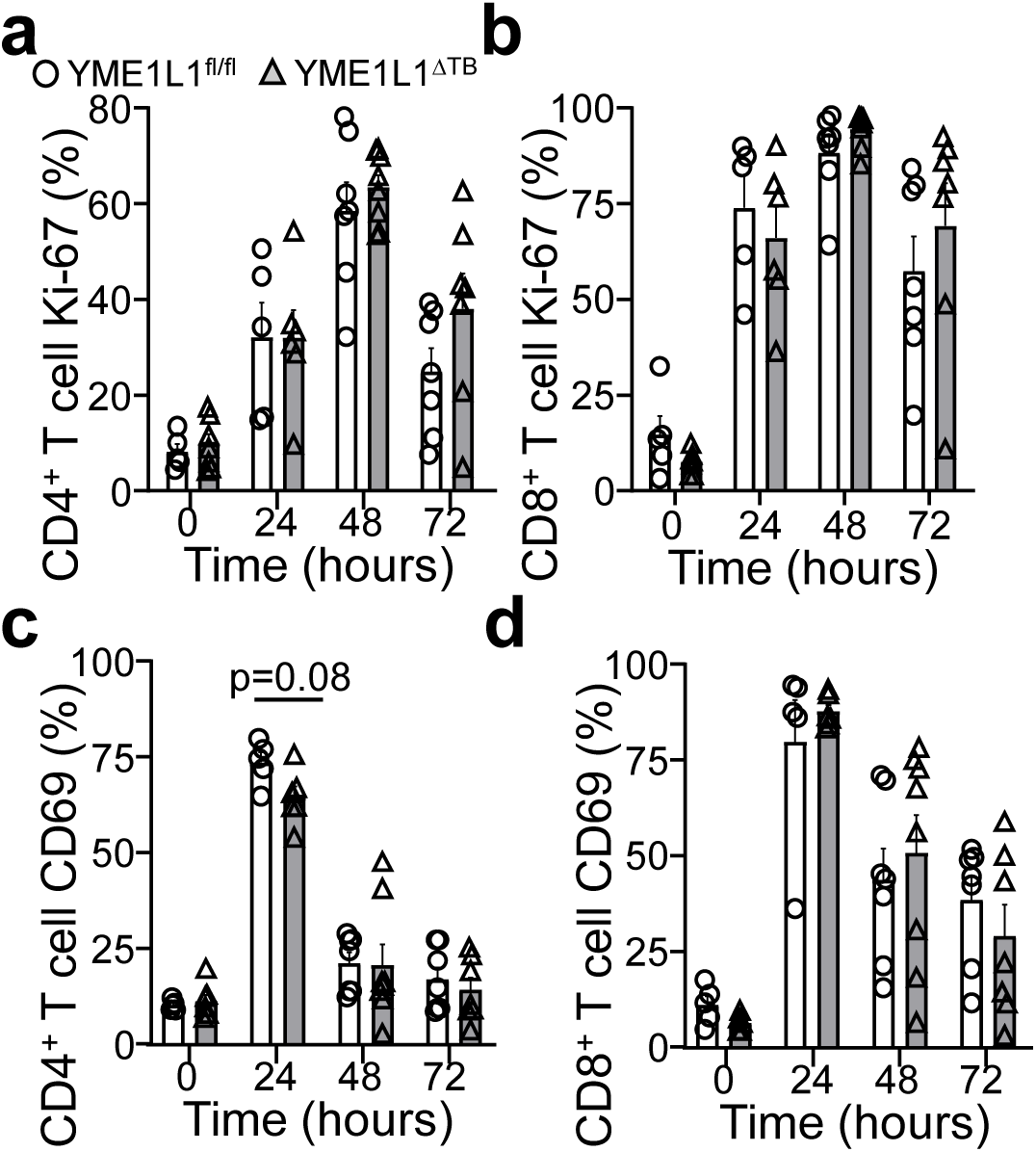
YME1L1-deficiency does not impact T cell proliferation and activation. CD4 and CD8 T cells were harvested from YME1L1^f/f^ littermate controls and YME1L1^ΔTB^ animals at steady state or upon indicated time after intraperitoneal administration of anti-CD3ε. **a-b**) Ki-67 staining in a) CD4 T cells, and b) CD8 T cells. **c-d**) Cd69 staining in c) CD4 T cells, and d) CD8 T cells (n=5-7). Statistical analysis: Mann-Whitney tests.

### Mitochondrial mass and function show minor alterations in the absence of YME1L1

Although the absence of YME1L1 in lymphocytes did not result in clear αβ T cell phenotype, this does not exclude a role for YME1L1 in mitochondrial biology in these cells. Therefore, we analysed the mitochondrial mass using Mitotracker Green (MTG), membrane potential using Tetramethylrhodamine Ethyl Ester (TMRE), and mitochondrial reactive oxygen species (ROS) production using MitoSOX Red (MSR) in the absence or presence of YME1L1 and upon T cell activation over three days. At steady state the three main lymphocyte populations, B, CD4 and CD8 T cells, showed a similar staining profile for MTG. Throughout the stages of activation up to 72 hours after anti-CD3ε administration, MTG staining intensity remained similar independently of the presence of YME1L1 (Fig. 6a-c).

**Figure 6.**
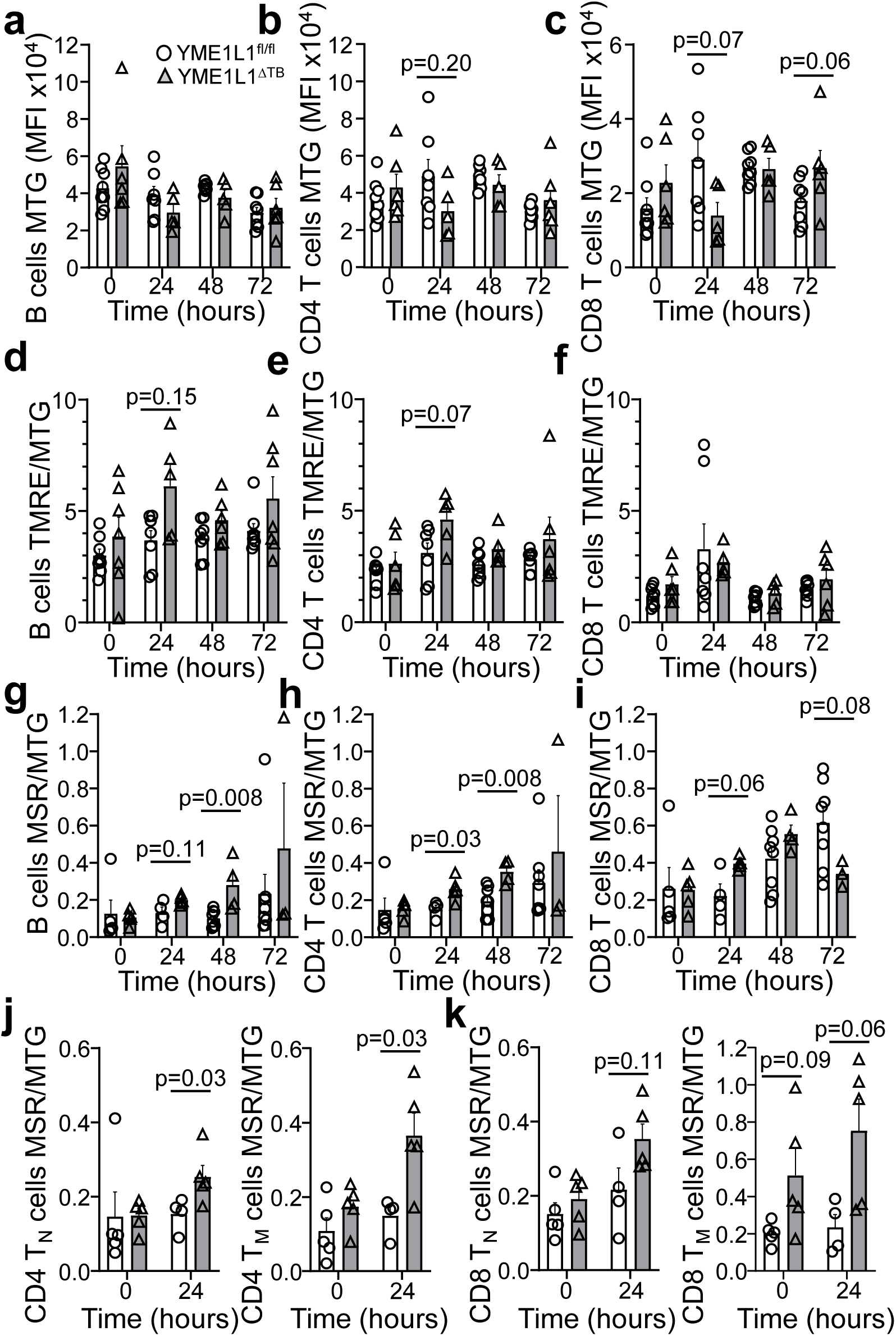
YME1L1-deficiency has limited impact mitochondrial dyes for mass, membrane potential and reactive oxygen production. CD4 and CD8 T cells or B cells were harvested from YME1L1^f/f^ littermate controls and YME1L1^ΔTB^ animals at steady state or upon indicated time after intraperitoneal administration of anti-CD3ε. **a-c**) Mean fluorescence intensity (MFI) of MitoTracker Green (MTG) in a) B cells, b) CD4 T cells, and c) CD8 T cells. **d-f**) Ratio of Tetramethylrhodamine, Ethyl Ester, Perchlorate (TMRE) over MTG staining in d) B cells, e) CD4 T cells, and f) CD8 T cells. **g-k**) Ratio of MitoSOX Red (MSR) over MTG staining in g) B cells, h) CD4 T cells, and i) CD8 T cells; j) naïve and memory CD4 T cells, and k) naïve and memory CD8 T cells (n=6-7). Statistical analysis: Mann-Whitney tests.

Similarly, mitochondrial membrane potential, measured by TMRE and normalised to mitochondrial mass, showed no difference at steady state between both genotypes (Fig. 6d–f). At 24 hours post-activation, B and CD4 T cells lacking YME1L1 displayed a trend towards increased TMRE staining compared with controls, an effect not observed in CD8 T cells (Fig. 6d–f). ROS production followed a similar pattern. At baseline, MSR staining was indistinguishable between groups. Upon activation, MSR signal increased in all lymphocytes over time, with YME1L1-deficient lymphocytes, especially B and CD4 T cells, showing a trend towards elevated ROS levels at 24 and 48 hours. Differences in CD8 T cells did not reach significance, with an indication that ROS levels may rapidly reduce at 72 hours post activation in the absence of YME1L1 (Fig. 6g–i).

Because circulating CD4 and CD8 T cells comprise both naïve and memory subsets, which may mask subtle differences, we next analysed these populations separately at baseline and 24 hours post-activation, when CD44 expression still reliably distinguishes previously naïve from previously activated T cells. This revealed that both naïve and memory CD4 T cells lacking YME1L1 exhibited increased MSR staining at 24 hours, whereas CD8 T cell subsets showed only a non-significant trend in the same direction (Fig. 6j–k).

### YME1L1-deficiency does alter mitochondrial shape in T cells

Our data shows that YME1L1 protein levels increase upon T cell activation, but the absence of YME1L1 in lymphocytes by itself does not markedly affect their development or levels and proportions in the spleen, despite small differences observed in γδ T cells, particularly their ability to make IFNγ or IL-17 [37]. Furthermore, T cell proliferation and activation, based on Ki-67 and CD69 expression, remained unaffected. Changes in mitochondrial mass, membrane potential and ROS production showed a trend towards reduced mitochondrial mass and an increase in ROS production in the absence of YME1L1, which may indicate possible compensatory mechanisms. In addition, this would suggest that mitochondrial shape, structure and size may still be altered, as previously reported in other cell types [11, 17, 23, 27–32].

To assess the consequence of the absence of YME1L1 in T lymphocytes, we investigated the proteolytic cleavage of OPA1, which in mice, depending on the tissue [38], can show up to five forms; two L-OPA1 and three S-OPA1 [11]. YME1L1-deficiency results in the absence of the first L-OPA1 processed form in T lymphocytes. In agreement with previous results in other cell types [17, 39], YME1L1-deficiency in T cells alters OPA1 cleavage (Fig.7a).

We then investigated the size and shape of mitochondria using electron microscopy (EM) images taken from memory CD8 T cells, the subset of T cells most studied with respect to their mitochondrial biology. For this, we established a quantitative image analysis workflow, as briefly explained in the Materials and Methods section. Mitochondria were automatically segmented using micro_SAM [40], implemented in Napar, open-source tools that can use deep learning models trained for identifying organelles in EM images. The variance in contrast across different EM images was considered, contrast adjusted, and a Fire Look up Table (LUT) applied to aid visualization.

Using this workflow, we screened several representative EM images (Fig. 7b). Although mitochondrial size cannot accurately be assessed in individual two-dimensional sections, we screened over 30 randomly imaged mitochondria from each genotype. The absence of YME1L1 in CD8 memory T cells did not alter the mitochondrial section area, nor their roundness compared to controls (Fig. 7c, d). However, assessment of cristae branching revealed an altered organisation in the absence of YME1L1 by a significant increase in branching (Fig. 7e). The size of cristae was assessed via the Feret diameter, and the distance between cristae assessed by automatically measuring the perpendicular distance between two parallel lines that are tangent to the extreme points of the shape, which revealed altered spacing between cristae in the absence of YME1L1 protein (Fig. 7f). This was confirmed by assessing the cristae tortuosity, i.e. the ratio between the length of the cristae to the straight-line distance between the two farthest extremities, in the both the presence and absence of YME1L1. In the absence of YME1L1, the cristae in T cells were shorter, less straight and more convoluted (Fig. 7g, Suppl. Fig. 1a-b).

**Figure 7.**
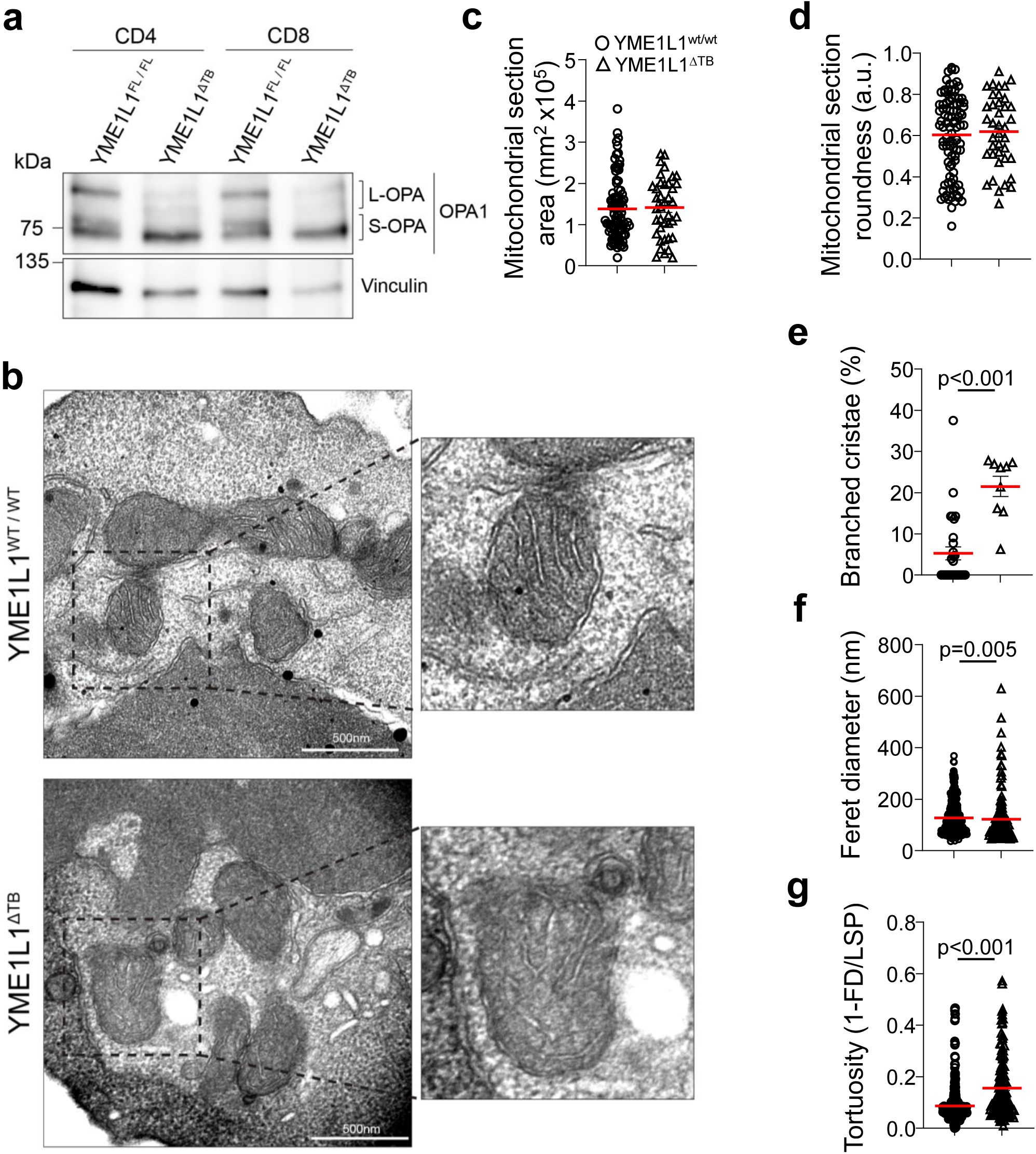
YME1L1-deficiency does impact mitochondrial fragmentation and shape in T lymphocytes. **a**) CD4 or CD8 T cells were enriched using AutoMACS selection, lysed and OPA1 presence determined. Shown is a representative Western blot from three biological repeats. **b-e**) Memory CD8 T cells were isolated from spleens from YME1L1^f/f^ littermate controls and YME1L1^ΔTB^ animals, purified via flow sorting (CD8^+^CD44^hi^), and analysed using an electron microscope. b) Representative EM images, were processed to segment mitochondria and cristae respectively, and measurements were made. c) Mitochondrial section area (nm2), d) Mitochondrial section roundness (4π·Area/Perimeter² in arbitrary units (a.u.)) (n=94 controls, n=37 YME1L1^ΔTB^), e) Percentage of branched cristae (n=29 controls, n=9 YME1L1^ΔTB^), f) Cristae Feret diameter (nm), and g) Tortuosity calculated as 1-Feret diameter (FD)/longest shortest path (LSP) (n=371 controls, n=138 YME1L1^ΔTB^). Statistical analysis: Mann-Whitney tests.

Collectively, these data show that in agreement with other cell types, YME1L1 has a role in cleaving OPA1, thereby influencing mitochondrial cristae organisation and shape in T lymphocytes.

## Discussion

We set out to determine whether the mitochondrial protease YME1L1 plays an important role in T cell biology, especially during activation where its levels are increased, given its established importance in mitochondrial dynamics and homeostasis in multiple other cell types. Contrary to expectations, T cell development and peripheral homeostasis were unaffected by YME1L1 deficiency. Furthermore, T cell activation and proliferation in response to TCR stimulation proceeded normally, and mitochondrial mass, membrane potential, and ROS production were largely unchanged, with only modest and transient differences. These results suggest that YME1L1, while critically regulating OPA1 and being upregulated upon T cell activation, is not essential for the fundamental processes driving T cell activation and proliferation.

These findings contrast with reports in fibroblasts, oocytes, neurons and cancer cells, where YME1L1 loss results in marked mitochondrial fragmentation, reduced membrane potential, and impaired cell viability [11, 17, 23, 25, 26, 28, 29]. The absence of major functional defects in T lymphocytes suggests that either T cells possess redundant or compensatory mechanisms, or that the demands placed on mitochondria during T cell activation differ from those in metabolically distinct cell types. It is possible that mitochondrial shaping proteins may buffer the loss of YME1L1, even though OPA1 cleavage remains altered.

We did detect subtle mitochondrial alterations in YME1L1-deficient T cells compared to their controls, including cristae branching and higher cristae tortuosity, in opposition with a role in fine-tuning cristae architecture. These changes did not translate into overt functional impairment under the experimental conditions tested, but they may become relevant under conditions of metabolic stress, infection, or chronic antigen stimulation, when possibly more of bioenergetic requirements of T cells are needed. In this regard, the modest shift observed in γδ T cell cytokine bias, with increased IFN-γ production and reduced IL-17 production, hints at context-specific roles of YME1L1 in shaping T cell effector functions.

Our work has shortcomings. We determined mitochondrial shape using two-dimensional images, which can hide additional features. However, using representative images and unbiased analysis of basic features we obtained a robust impression of the cristae structure in the absence of YME1L1 in memory CD8 T cells. Furthermore, we used group sizes of 6 mice each to avoid emphasis on modest changes. These changes may be small but can be relevant for some functions of T cells. The functional relevance was tested only with basic parameters, the presence of Ki67 and the expression of CD69. Although these showed no differences between the cells with or without YME1L1, additional functional differences cannot be excluded. Lastly, we determined levels and proportions of B cells but relied on bystander activation caused by T cell activation for an initial assessment of their biology in the absence of YME1L1. This does not exclude a specialised role in some of the B cell functions. More detailed work would require the extension into infection or immunisation models and lymphocyte differentiation, for which we have no supporting data of gaining potential new insights.

Our work demonstrates that YME1L1 is dispensable for core aspects of T cell development and acute activation yet subtly affects mitochondrial ultrastructure and γδ T cell effector programming. These findings highlight the importance of testing mitochondrial regulators in a cell-type–specific context and underscore that negative data are essential for refining our understanding of the complexity and redundancy in mitochondrial quality control pathways.

## Materials and methods

### Mice

*Yme1l*^fl/fl^ mice were kindly provided by Dr. Thomas Langer [17], Rag1^Cre^ was kindly provided by Dr. Terence Rabbitts [34]. Mice were bred at the Gulbenkian Institute for Molecular Medicine, Lisbon, Portugal. Male and female mice, aged and sex-matched, at 8-12 weeks of age were used. Animals were housed in IVC cages with temperature-controlled conditions under a 12-h light/dark cycle with free access to drinking water and food. All mice were kept in specific pathogen-free conditions. All mice were stringently genotyped by PCR. All animal experimentation complied with regulations and guidelines of, and was approved by, the Direção-Geral de Alimentação e Veterinária Portugal and the local ethical review committee (Orbea).

### Flow cytometry

Flow cytometry experiments were performed at the flowcytometry platform of the Gulbenkian Institute for Molecular Medicine. Single-cell suspensions from spleen were made after dissection and meshing the spleen through a 40µm filter, folowed by blood cells (RBC) lysis (150mM NH4Cl, 10mM KHCO3, 0.1mM Na2EDTA) at RT for 5 minutes. Cells were subsequently stained with antibodies (Supplementary Table 1), according to the agreed standards [40]. Samples were run on a Fortessa X20 cytometer (BD Biosciences) or Aurora (Cytek) and analysed with FlowJo X software (TreeStar).

For mitochondrial dye staining, cells were incubated in incomplete Incove’s modified Dulbeco’s Medium (IMDM) with 5% Glutamine and stained with 10 nM MitoTracker Green (MTG), 5nM tetramethylrhodamine ethyl ester (TMRE) and 2.5 µM MitoSox Red (MSR) (all Thermo Fisher Scientific) at 37°C. MTG and TMRE were stained for 30 min and MSR was stained for 10 min. All before extracellular Ab staining.

### Western Blots

Cell suspensions from spleen and lymph nodes were made after dissection and meshing through a 40µm filter, followed by RBC lysis. Cell suspensions were incubated with CD4-, CD8- or CD19-APC antibodies (Supplementary Table 1) and subsequently with anti-APC Microbeads (Milteny Biotech), for the isolation of CD4+ or CD8+ T cells or CD19+ B cells, respectively, by the autoMACS Pro Separator (Milteny Biotech). Cell lysis was performed in Triton lysis buffer (1% Triton X-100, 1mM ETDA, 50mM Tris/HCl pH 8.0, 150mM NaCl and freshly added complete EDTA free protease inhibitor) on ice for 1hr. The samples were centrifuged at 17000 g at 4 °C for 15 min and the supernatant was collected. Protein quantification was performed using Bradford protein assay (Bio-Rad) according to the manufacturer’s protocol and the samples were denatured at 95°C for 5 min in 6x Laemmli Buffer (50mM Tris-HCL pH 6.8, 60% glycerol, 4% SDS, 0,1% bromophenol blue and freshly added 5% 2-mercaptoethanol). 15µg, unless otherwise stated, of protein were loaded per well in 12% acrylamide/bis-acrylamide SDS PAGE gels, varying acrylamide/bis-acrylamide percentages as necessary, blotted in PVDF membranes, which were blocked with 5% non-fat dry milk in 1x TBS (150mM NaCl, 20mM Tris, 0.1% Tween-20, pH 7.6) for 1 h at RT. The membranes were incubated with each primary antibody (Supplementary Table 1) overnight at 4 °C, washed 3 times in TBS, followed by 1hr incubation with the respective HRP-conjugated secondary antibodies in 5% non-fat milk in 1x TBS at RT. After 3 times washing with 1x TBS, the membranes were developed with chemiluminescent detection using Clarity^TM^ Western ECL Substrate (BioRad), in an Amersham ImageQuant 800 (Cytiva).

For OPA1 western blotting, the same steps were performed with the exception of the cell lysis, which was performed in RIPA Buffer (50mM Tris/HCl pH 8.0, 1mM EDTA, 1% Triton X-100, 0.5% sodium deoxycholate, 0.1% SDS, 150mM NaCl) for 1hr on ice followed by sonication for 30 cycles of 15sec on/off.

### EM and analysis

Microscopy experiments were performed at the Electron Microscopy and Bioimaging facilities of the Gulbenkian Institute for Molecular Medicine. After cell sorting, cells were left to rest for 1 hour in IMDM. Cells were seeded in 24 well plates with 0.01mg/mL Poly-D-Lysine (PDL) coated cover slips (13mm) and centrifuged at 200 xg for 10 min to aid attachment. Cell seeding density was from 250K to 1M cells depending on the population. After this, the supernatant was removed and cells were fixed with 2% formaldehyde, 2.5% glutaraldehyde in 0.1M phosphate buffer pH7.4 for 1h at RT. Cells are then washed in 0.1M phosphate buffer (PB). Postfix was done in 1% Osmium, 1.5% potassium ferrocyanide for 30 min, on ice, in the dark and washed 2 times with 0.1M PB and once in dH2O. Cells were stained with 1% tannic acid for 20 minutes on ice and washed 5x with dH2O. Cells were stained with 0.5% uranyl acetate in dH2O for one hour at RT, in the dark. Samples were dehydrated with a series of increasing ethanol concentrations (50%, 70%, 90%, 3×100%). Infiltrating/embedding was made in Epon resin (EMbed 812). After polymerising resin at 60 LJC overnight, glass was removed by immersion in liquid nitrogen and resin blocks were sectioned *en face* at 70nm thickness using a UC7 ultramicrotome (Leica) and a diamond knife (Diatome). Sections were collected on copper mesh grids and post-stained with 2% uranyl acetate in 70% methanol followed by Reynold’s lead citrate. Samples were imaged with a FEl Tecnai G2 Spirit BioTWIN operating at 120 keV and equipped with an Olympus-SIS Veleta CCD Camera. Whole cell images were acquired at a magnification of 11.5kx and mitochondria were acquired at 43kx. All reagents and materials were purchased from Electron Microscopy Sciences unless otherwise stated.

Roundness was determined as 4 x area/(π x major_axis^2^), or the inverse of the aspect ratio. Mitochondria were automatically segmented using a deep learning model trained for identifying organelles in EM images, followed by a re-segmentation using a trained pixel-classifier, to distinguish matrix from cristae within mitochondria (Supplementary Figure 1). After having segmented separately whole mitochondrial cross-sections and their matrix vs cristae we developed a set of custom-made MACROS in FIJI to automatically measure skeletal length and Feret length of cristae, and a derived measure of “tortuosity”, as well as the inter-cristae distance. The macros are presented in Supplementary material and can be obtained from the authors upon request.

### Statistical analysis

Statistical analyses were performed using GraphPad Prism Software (GraphPad Prism version 9 for Windows, GraphPad Software, San Diego, California USA), details described in each figure legend where applicable. n denotes the total number of independent biological samples (mice). Error bars represent S.E.M. Mann–Whitney analysis was applied to compare ranks between two groups with a p value of < 0.05 considered significant.

## Supporting information

Macros used

## Acknowledgements

We would like to thank the technical support of the Gulbenkian Institute for Molecular Medicine platforms for flow cytometry, rodent, histology, electron microscopy and bioimaging. We are grateful for additional bioimaging analysis assistance from Alexandre Lopes. The project that gave rise to these results has received funding from the following sources: ”la Caixa” Foundation under the grant agreement LCF/PR/HR24/52440020; FCT – Portuguese Foundation for Science and Technology under grant agreements (UI/BD/151173/2021) (G.M.), PPBI-POCI-01-0145-FEDER-022122, and V.A.M. was an FCT researcher (IF/01693/2014; IMM/CT/27-2020; 2021.03613.CEECIND).

## Author contributions

G.M. performed the phenotyping of the mice using flow cytometry. M.J. performed the biochemical assays to determine protein contents. R.M. performed the proliferation assays, mitochondrial dye intensity assays using flow cytometry, and performed the analysis of the EM data. J.F. performed the genotyping and colony maintenance, ordering of stock and helped with the experiments. G.M. helped design and implement the mitochondrial imaging analysis, collected by M.J.H. V.M. and M.V. coordinated the work, secured funding and wrote the manuscript. All authors contributed to the writing of the manuscript.

## Data availability statement

All data will be made available upon reasonable request.

