## Supplemental table 1 for "YME1L1 is Dispensable for T Lymphocyte Activation Despite its Upregulation and Activity"

**Supplement table 1.** Antibody list and details of those used for flow cytometry and Western blots.

| **Marker** | **Fluorophore** | **Isotype** | **Reactivity** | **Clone** | **Manufacturer** |
|---|---|---|---|---|---|
| Live/Dead near-IR fluorescent-reactive dye | "APC/Cy7" | | | | Invitrogen |
| CD4 | APC | Rat IgG2a, k | mouse | RM4-5 | Biolegend |
| CD8 | APC | Rat IgG2a, k | mouse | 53-6.7 | Biolegend |
| CD19 | APC | Rat IgG2a, k | mouse | 1D3/CD19 | Biolegend |
| CD4 | PerCP/Cy5.5 | Rat IgG2a, k | mouse | RM4-5 | Biolegend |
| CD8a | Super Bright 600 | Rat IgG2a, k | mouse | 53-6.7 | Invitrogen |
| TCRb | Pacific Blue | Armenian hamster IgG | mouse | H57-597 | Biolegend |
| TCRg/d | FITC | Armenian hamster IgG | mouse | UC7-13D5 | Biolegend |
| CD44 | Alexa Fluor 700 | Rat IgG2b, k | human/mouse | IM7 | Invitrogen |
| CD25 | Brilliant Violet 785 | Rat IgG1, λ | mouse | PC61 | Biolegend |
| CD27 | APC | Armenian hamster IgG | mouse/human | LG.3A10 | Biolegend |
| CD69 | PE/Cy7 | Armenian hamster IgG | mouse | H1.2F3 | Biolegend |
| CD24 | Brilliant Violet 510 | Rat IgG2b, k | mouse | M1/69 | Biolegend |
| CD19 | Pacific Blue | Rat IgG2a, k | mouse | 6D5 | Biolegend |
| CD62L | Brilliant Violet 510 | Rat IgG2a, k | mouse | MEL-14 | Biolegend |
| CD127 | PE/Cy7 | Rat IgG2a, k | mouse | A7R34 | Biolegend |
| CD4 | PE/Cyanine7 | Rat IgG2a, k | mouse | RM4-5 | Biolegend |
| CD45.2 | APC | Mouse (SJL) IgG2a, k | mouse | 104 | Biolegend |

| | | | | | |
|---|---|---|---|---|---|
| **Marker** | **Fluorophore** | **Isotype** | **Reactivity** | **Cat #** | **Manufacturer** |
| YME1L1 | | Rabbit | human/mouse | 1150-1-AP | Proteintech |
| b-actin | | Rabbit | human/mouse | #49675 | Cell Signaling |
| Vinculin | | Mouse | human/mouse | V9131 | Sigma-Aldrich |
| OPA1 | | Mouse | human/mouse | 612607 | BD Biosciences |
