## Supplementary material for "YME1L1 is Dispensable for T Lymphocyte Activation Despite its Upregulation and Activity": Macros used

**Supplement Macros**

1. **FIJI MACRO for mitochondrial number; section area, section roundness, section thickness, cristae thickness, number and junction number.**

```
// open the raw data and the segmented, then click on the segmented image to proceed (the raw is just for checking if results make sense)
// the segmented image was produced in Ilastik and gray = 1 for backg, =2 for mito matrix and =3 for cristae

// Count manually the mitos and score them in the table
// then proceed to skeletonize and measure the cristae

run("Properties...", "channels=1 slices=1 frames=1 pixel_width=1.07 pixel_height=1.07 voxel_depth=80 global");
run("Set Measurements...", "area shape feret's display redirect=None decimal=2");

// user selects the image of the segmentation of the mitos
run("Enhance Contrast", "saturated=0.35");
run("Median...", "radius=7");
run("Duplicate...", "title=mito");
run("Duplicate...", "title=matrix");
run("Duplicate...", "title=cristae");

selectImage("mito");
setThreshold(2, 3, "raw");
run("Convert to Mask");
selectImage("matrix");
setThreshold(2, 2, "raw");
run("Convert to Mask");
selectImage("cristae");
setThreshold(3, 3, "raw");
run("Convert to Mask");

run("Tile"); //tidy up the screen

// measure mito sections
selectImage("mito");
run("Analyze Particles...", "size=100-Infinity display exclude clear include");
selectWindow("Results");
run("Summarize");

waitForUser("Mito section shape", "Record mean area + round, then OK");
close("Results");
selectImage("mito");
run("Local Thickness (masked, calibrated, silent)");
run("Histogram", "bins=256 use x_min=0 x_max=500 y_max=Auto");
waitForUser("Mito section shape", "Record # mitos + mean & mode thicknes, then close histo and colored image, then click OK");
close("Histogram of mito_LocThk");
close("mito_LocThk");
```

```
// proceed to matrix measurements

selectImage("matrix");
run("Local Thickness (masked, calibrated, silent)");
run("Histogram", "bins=256 use x_min=0 x_max=500 y_max=Auto");
waitForUser("Matrix thickness", "Record mean + mode thickness, then close histo and click
OK to continue");
close("Histogram of matrix_LocThk");
close("matrix_LocThk");

// proceed to cristae thickness measurements

selectImage("cristae");
// "cristae" with less than 200nm^2 will NOT be measured after this!
run("Analyze Particles...", "size=200-Infinity show=Masks display exclude clear include
add");
selectWindow("Results");
run("Summarize");
waitForUser("Cristae Feret", "Record # cristae + mean cristae Feret, then click OK to
continue");
close("cristae");
selectImage("Mask of cristae");
run("Invert LUT");
rename("cleanCristae");
run("Duplicate...", "title=cristaeSkel");

selectImage("cleanCristae");
run("Local Thickness (masked, calibrated, silent)");
run("Histogram", "bins=256 use x_min=0 x_max=100 y_max=Auto");
waitForUser("Cristae thickness", "Record mean & mode thickness, then close histo and click
OK to continue");

// proceed to cristae skeleton measurements

selectImage("cristaeSkel");
run("Skeletonize (2D/3D)");
run("Analyze Skeleton (2D/3D)", "prune=none calculate show");
selectWindow("Results");
Table.sort("# Junctions");
run("Summarize");

waitForUser("END!", "Record #Junctions and Mean 'longest short path'");

// run("Merge Channels...", "c1=mito c2=matrix c3=cristae create");

// --- after your first macro finishes ---
waitForUser("Did you copy the value you needed? Click OK to continue.");

// Clean up before restarting
run("Close All");
roiManager("reset");
run("Clear Results");
```

### 2. FIJI MACRO for Cristae FD

```
macro "Run My Analysis Action [r]" {
   // your code here
// --- IMAGE PROCESSING ---
run("Enhance Contrast", "saturated=0.35");
run("Median...", "radius=7");
// Duplicate channels
run("Duplicate...", "title=mito");
run("Duplicate...", "title=matrix");
run("Duplicate...", "title=cristae");
// Threshold & convert to mask
selectImage("mito");
setThreshold(2, 3, "raw");
run("Convert to Mask");
selectImage("matrix");
setThreshold(2, 2, "raw");
run("Convert to Mask");
selectImage("cristae");
setThreshold(3, 3, "raw");
run("Convert to Mask");

selectImage("matrix");
close();
selectImage("mito");
close();

selectImage("cristae");
run("Analyze Particles...", "size=450-Infinity pixel circularity=0.00-0.50 show=Masks add
clear");
selectImage("Mask of cristae");
roiManager("deselect");
roiManager("save", "/Users/ricardomachado/Desktop/rois.zip");
run("Invert");

maxy = roiManager("count");
print("==========");
for (roiIndex = 0; roiIndex < maxy; roiIndex++) {

        roiManager("deselect");
        roiManager("delete");
        roiManager("open", "/Users/ricardomachado/Desktop/rois.zip");
        selectImage("Mask of cristae");
        // Select the ROI for this iteration
        roiManager("Select", roiIndex);
        run("Create Mask");
        setOption("BlackBackground", true);
        run("Skeletonize");
        selectImage("Mask");

        // This command will analyze the single, skeletonized object
        // and add its properties to the results table.
        run("Analyze Particles...", "size=1-Infinity clear add");
```

```
        // Now, get the result from the single row in the table.
        value = getResultString("Feret", 0);
        print("Feret of skeleton " + roiIndex + " " + value);

        // Clean up for the next loop
        close("Mask");
        run("Clear Results");
}

// Final cleanup
close("*");
roiManager("reset");
close("results");
}
```

3. **FIJI MACRO for Cristae LSP**

```
macro "Run My Analysis Action [r]" {
   // your code here
//open("C:/Users/alopes/Downloads/lixo/Ricardo/ILASTIK_KO_6.tiff");
   // your code here
// --- IMAGE PROCESSING ---
run("Enhance Contrast", "saturated=0.35");
run("Median...", "radius=7");
// Duplicate channels
run("Duplicate...", "title=mito");
run("Duplicate...", "title=matrix");
run("Duplicate...", "title=cristae");
// Threshold & convert to mask
selectImage("mito");
setThreshold(2, 3, "raw");
run("Convert to Mask");
selectImage("matrix");
setThreshold(2, 2, "raw");
run("Convert to Mask");
selectImage("cristae");
setThreshold(3, 3, "raw");
run("Convert to Mask");

selectImage("matrix");
close;
selectImage("mito");
close;

selectImage("cristae");
run("Analyze Particles...", "size=450-Infinity pixel circularity=0.00-0.50 show=Masks add
clear");
selectImage("Mask of cristae");
```

```
roiManager("deselect");
roiManager("save", "/Users/ricardomachado/Desktop/rois.zip");
run("Invert");

// Ask user which ROI to pick
//Dialog.create("Pick ROI");
//Dialog.addNumber("Enter ROI index:", 1);
//Dialog.show();
//roiIndex = Dialog.getNumber();

// Ask user which ROI to pick
//Dialog.create("Pick ROI");
//Dialog.addNumber("Enter ROI index:", 1);
//Dialog.show();
//roiIndex = Dialog.getNumber();

maxy = roiManager("count");
print("===========");
for (roiIndex = 0; roiIndex < maxy; roiIndex++) {

	roiManager("deselect");
	roiManager("delete");
	roiManager("open", "/Users/ricardomachado/Desktop/rois.zip");
	selectImage("Mask of cristae");
	// Select the ROI
	roiManager("Select", roiIndex);
	run("Create Mask");
	setOption("BlackBackground", true);
	run("Skeletonize");
	//selectImage("Mask");
	run("Analyze Skeleton (2D/3D)", "prune=none calculate show");
	selectImage("Tagged skeleton");
	close;
	selectImage("Longest shortest paths");
	close;
	selectImage("Mask");
	close;
	//waitForUser;

	value = getResultString("Longest Shortest Path", 0);
	print("Longest Shortest Path " + roiIndex + " " + value);


}


// --- after your first macro finishes ---
//waitForUser("Did you copy the value you needed? Click OK to continue.");

// Clean up before restarting
close("*");
roiManager("reset");
close("results");
close("Branch information");}
```
