## Supplemental figure 1 for "YME1L1 is Dispensable for T Lymphocyte Activation Despite its Upregulation and Activity"

### Supplementary Figure 1

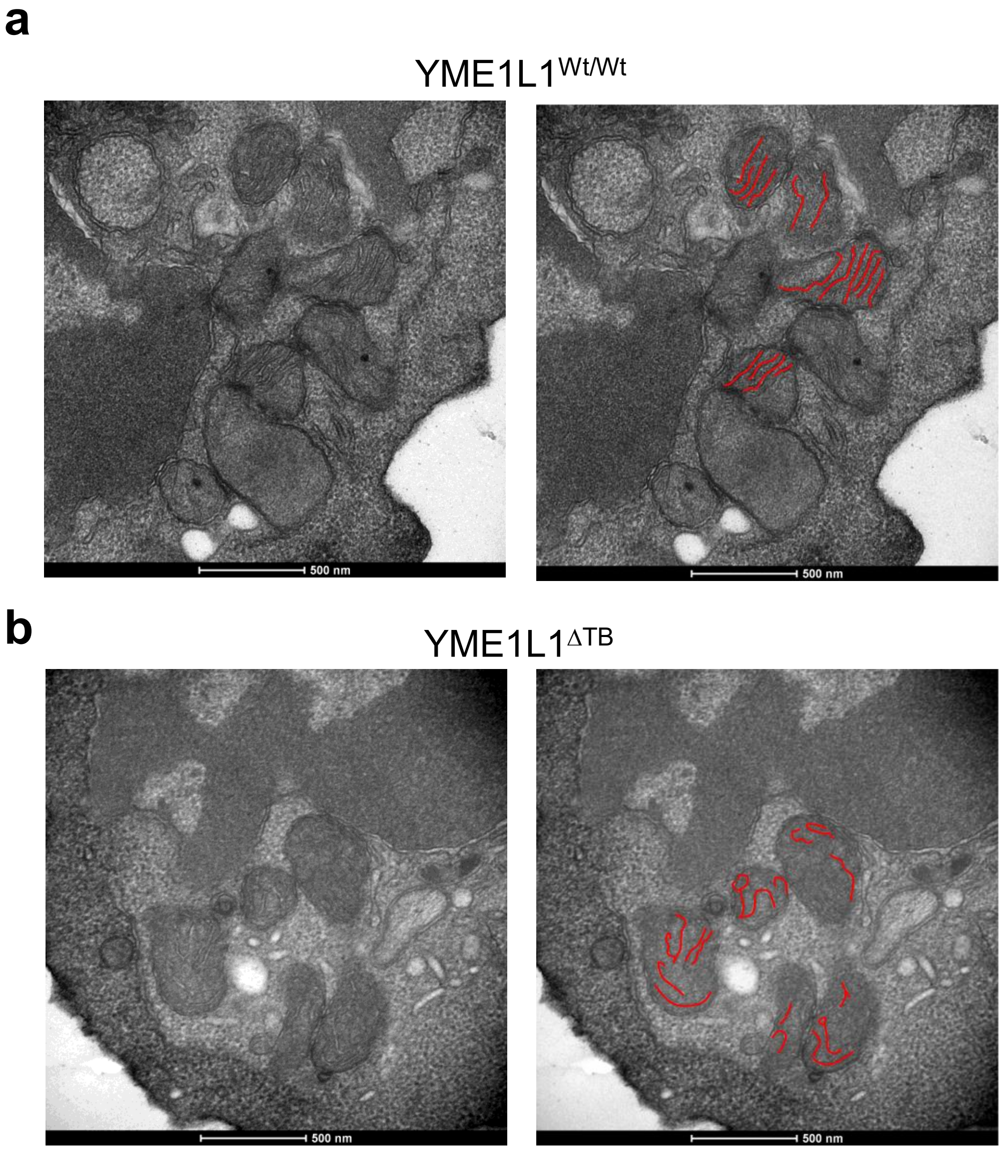


**Supplementary Figure 1. YME1L1-deficiency results in altered cristae morphology in T lymphocytes**. CD8 memory T cells were flow sorted based on CD8 and CD44. Representative transmission electron microscopy (TEM) images from **a**) YME1L1$^{Wt/Wt}$ and **b**) YME1L1$^{\Delta TB}$ mice. Left panels show unannotated images; right panels show the corresponding images with red lines indicating cristae shape. Analysis were performed on unannotated images. Scale bars are shown in each image.
